# Faster than the brain’s speed of light: Retinocortical interactions differ in high frequency activity when processing darks and lights

**DOI:** 10.1101/153551

**Authors:** Britta U. Westner, Sarang S. Dalal

## Abstract

Visual processing of dark visual stimuli has been hypothesized to occur faster relative to bright stimuli. We investigated the timing, processing, and propagation of neural activity in response to darks and lights, operationalized as light offset and onset, in the human visual system by recording electroretinography (ERG) simultaneously with magnetoencephalography (MEG) in humans. We discovered that dark onset resulted in 75–95 Hz retinal activity that we call the *dark retinal oscillatory potential*, occurring with the same latency as the analogous but more broadband (55–195 Hz) oscillatory potential at light onset. Both retinal oscillations coupled with subsequent cortical activity of corresponding bandwidths, but cortical responses for darks indeed occurred earlier than for lights. Darks therefore propagate from retina to cortex more quickly than lights, potentially resulting from a thalamic advantage. Furthermore, we found that this propagation is effectuated by high frequency retinocortical coupling of narrow bandwidth for darks but wide bandwidth for lights.

## Introduction

The visual system has long been known to segregate processing of light increments and light decrements into parallel ON and OFF pathways (Hartline, 1938). As these pathways involve different computations and neural structures, the relative speed with which information processed by these pathways propagate could very well vary between them. A behavioral advantage for the detection of dark objects or light decrements over bright objects has indeed been reported in several psychophysical studies (e.g., Blackwell (1946); Krauskopf (1980); Bowen et al. (1989); Chubb and Nam (2000); Buchner and Baumgartner (2007)). More recently, Komban et al. (2011) reported that the behavioral advantage for darks vanished in their experiment when the binary noise background was corrected for the irradiation illusion, a phenomenon in which light objects on a dark background seem larger than the opposite (Galilei, 1632; von Helmholtz, 1867).

These findings raise the question of at which stage of the visual system do such functional asymmetries in the ON and OFF pathways emerge. Numerous studies have investigated such asymmetries at various levels of the visual system and in several species.

In visual cortex, responses to light decrements are found to be stronger than responses to light increments in both electroencephalography (EEG) and functional magnetic resonance imaging (fMRI) recordings (Zemon et al., 1988; Zemon et al., 1995; Olman et al., 2008). Multiunit recordings from cat visual cortex show faster response latencies (defined as 40% of maximum response) in OFF-dominated cortical sites (Komban et al., 2014). Those functional results could be explained by studies showing neuronal differences between the ON and OFF pathways in visual cortex: the number of geniculate afferents at the representation of the area centralis in cat visual cortex was shown to be higher in the OFF pathway (Jin et al., 2008). Whether advantages for the processing of darks arise at the cortical or thalamic level is not clear: Yeh et al. (2009), for example, reported more black-dominant neurons in layers 2 and 3 of primary visual cortex (V1) of macaque monkeys, but a balanced amount of black- and white-dominant neurons in the thalamic input layer 4c of visual cortex. On the other hand, Xing et al. (2010) showed temporal advantages for dark stimuli in the thalamus input layer 4c of visual cortex, but not in the later stages of cortical visual processing. A potential advantage for dark stimuli at the thalamic level is further supported by a study of Jin et al. (2011), which reported faster processing for light decrements than increments in the lateral geniculate nucleus (LGN) of the cat thalamus.

At the retinal stage, evidence for functional asymmetries in the ON and OFF pathways is mixed. While some studies find no asymmetries at all (Kremers et al., 1993; Benardete and Kaplan, 1997; Benardete and Kaplan, 1999), others do report differences in ON and OFF processing. Several studies show faster responses for light decrements in the retina (Copenhagen et al., 1983; Zaghloul et al., 2003; Burkhardt et al., 2007; Gollisch and Meister, 2008; Nichols et al., 2013). However, Chichilnisky and Kalmar (2002) reported this temporal advantage only for the initial response, whereas the time to peak was shorter for ON but not OFF ganglion cells (also see Lankheet et al. (1998)). It has been hypothesized that OFF bipolar cells are faster in their response kinetics, since no biochemical sign inversion of the light response is needed - in contrast to ON bipolar cells (Nawy and Jahr, 1990; Chichilnisky and Kalmar, 2002). Furthermore, numerous studies across different species suggest that more neuronal resources are allocated to the OFF pathway (Balasubramanian and Sterling, 2009): for example, there are twice as many OFF than ON diffuse bipolar cells in the fovea of the macaque retina (Ahmad et al., 2003), OFF ganglion cells have narrower dendritic and thus narrower receptive fields than their ON counterparts (Wässle et al., 1981; Morigiwa et al., 1989; Dacey and Petersen, 1992; DeVries and Baylor, 1997) and show more overlap than ON dendritic fields (Borghuis et al., 2008). However, it has also been shown that OFF cell currents are rectified by ON cells (Zaghloul et al., 2003; Liang and Freed, 2010).

Our study focuses on the potential functional differences following flash onsets and offsets in the visual system to address the question whether darks are indeed processed faster than lights. By simultaneously recording the retinal and cortical responses, we aim at mapping the shape and timing of activity patterns at different stages of the human visual system to gain functional evidence at which stage such differences may arise.

### Retinal potentials and high frequency oscillations

Retinal potentials in response to full-field flashes have been used in the clinical assessment of retinal function for some decades (Marmor et al., 1989; Marmor et al., 2009) and are therefore well described. These potentials, which are seen in the electroretinogram (ERG), reflect the summed activity of the retinal network and arise from different processing stages (Frishman, 2013). While light onset is followed by several potentials, most importantly the negative, photoreceptor-generated a-wave (Perlman, 2001; Frishman, 2013) and the positive b-wave, which is mostly driven by ON bipolar cells (Sieving et al., 1994; Frishman, 2013; Vukmanic et al., 2014), light offset is followed by one positive deflection, the d-wave, which has its origin in several pathways, amongst them the cone photoreceptors and OFF pathways (Sieving et al., 1994; Perlman, 2001; Ueno et al., 2006; Frishman, 2013). The retina’s response to flash onset also comprises an onset-locked high frequency activity, the so-called oscillatory potential, first described by (Fröhlich, 1914). It involves frequencies centered around 120Hz and up to 200Hz (Kozak, 1971; Munk and Neuenschwander, 2000; Todorov et al., 2016). The precise mechanisms and the cellular origin of the oscillatory potential are still unclear; an involvement of ganglion, amacrine, and bipolar cells, possibly in a feedback loop, is discussed (Doty and Kimura, 1963; Perlman, 2001; Kenyon et al., 2003; Frishman, 2013). Kozak (1971) describes a similar but slower oscillation (65–100 Hz) in response to light offset in cats; whether there is an analogous high frequency oscillation following light offset in humans has not previously been demonstrated.

There is evidence that the retinal oscillatory potential is directly transmitted to visual cortex (Lopez and Sannita, 1997; Castelo-Branco et al., 1998; Sannita et al., 1999; Heinrich and Bach, 2001; Neuenschwander et al., 2002; Todorov et al., 2016). Other studies, however, have suggested that retinal and cortical high frequency activity are two distinct processes (Doty and Kimura, 1963; Molotchnikoff et al., 1975; Heinrich and Bach, 2004).

In our study, we use retinal and cortical high frequency activity to investigate whether darks are processed faster than lights across these different stages of the human visual system. The simultaneous recording of retinal and cortical activity furthermore enables a detailed comparison of the temporal dynamics and oscillatory patterns regarding retinocortical coupling and the transmission of the oscillatory potential to visual cortex.

We recorded the retinal and cortical responses to light flash onsets and offsets of 250 full-field light flashes with a duration of 480 ms. Retinal and cortical activity was measured simultaneously in 10 healthy participants using ERG and magneten-cephalography (MEG), thereby enabling a direct comparison of retinal and cortical high frequency activity and its propagation through the visual system.

## Results

### Retinal evoked potentials

Retinal activity was measured using disposable Dawson-Trick-Litzkow (DTL) fiber electrodes. The across-subjects average of retinal activity following light onset revealed the characteristic ERG flash response (Figure 1A). The negative-going a-wave had an average latency of 24.2 ms across subjects, followed by the positive-going b-wave at 79.9 ms. The data also shows the c-wave, which is generally seen with longer flashes (Hanitzsch et al., 1966; Skoog and Nilsson, 1974), but of no further interest in this study. Figure 1B shows the retinal response to light offset with the d-wave peaking at 25.2 ms after light offset (*n* = 8: for two subjects the peak was not identifiable). Latencies were compared using Bayesian Paired Samples T-Tests across participants. Results from these tests are reported as Bayes Factors (BF), which express the support of the alternative hypothesis over the null hypothesis (also see Method details, section Statistical testing). Following a categorization by Jeffreys (1961), *BF* > 3 are considered moderate evidence, *BF* > 10 strong evidence, and *BF* > 30 very strong evidence for the alternative hypothesis model (Lee and Wagenmaker, 2014). The comparison of d-wave and b-wave latencies yielded very strong evidence for a difference across participants (Bayes Factor *BF* = 90.42), with the d-wave being faster. There was no support for a peak latency difference between the a-wave and the d-wave: *BF* = 0.649.

**Figure 1:**
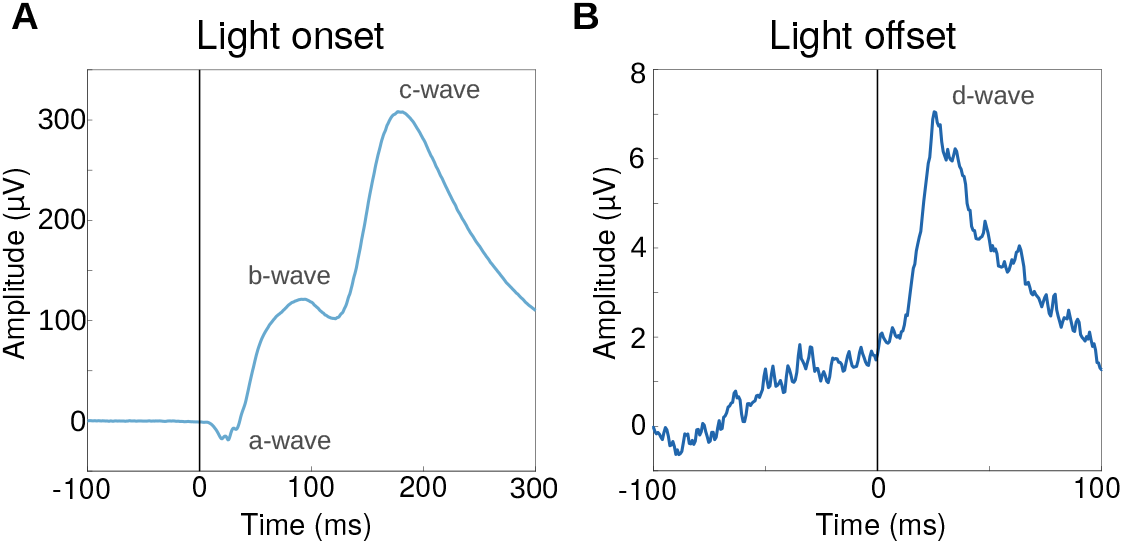
Retinal evoked potentials. **A** Retinal potentials following flash onset, averaged across subjects. **B** ERG response to light offset, averaged across subjects.

### Retinal and cortical high frequency activity

As illustrated in Figure 2A, high-pass filtering the light onset ERG data at 55 Hz reveals a high frequency burst. This oscillatory potential has high fidelity across trials and even individuals and is thus clearly represented in the group average. Figure 3 shows the quality of the high frequency responses from the retina and cortex of a representative subject. A comparable pattern in the light-off data is only evident when looking at a narrow frequency band of 75-95 Hz (Figure 2B). This is supported by the fact that across subjects, only this frequency band shows a significant increase in intertrial coherence (ITC) after light offset compared to baseline (Figure 2C). We call this phase-consistent high frequency activity the *dark retinal oscillatory potential*, as a novel darkness-evoked analogue to the classic light-evoked *oscillatory potential* exhibited by the retina.

**Figure 2:**
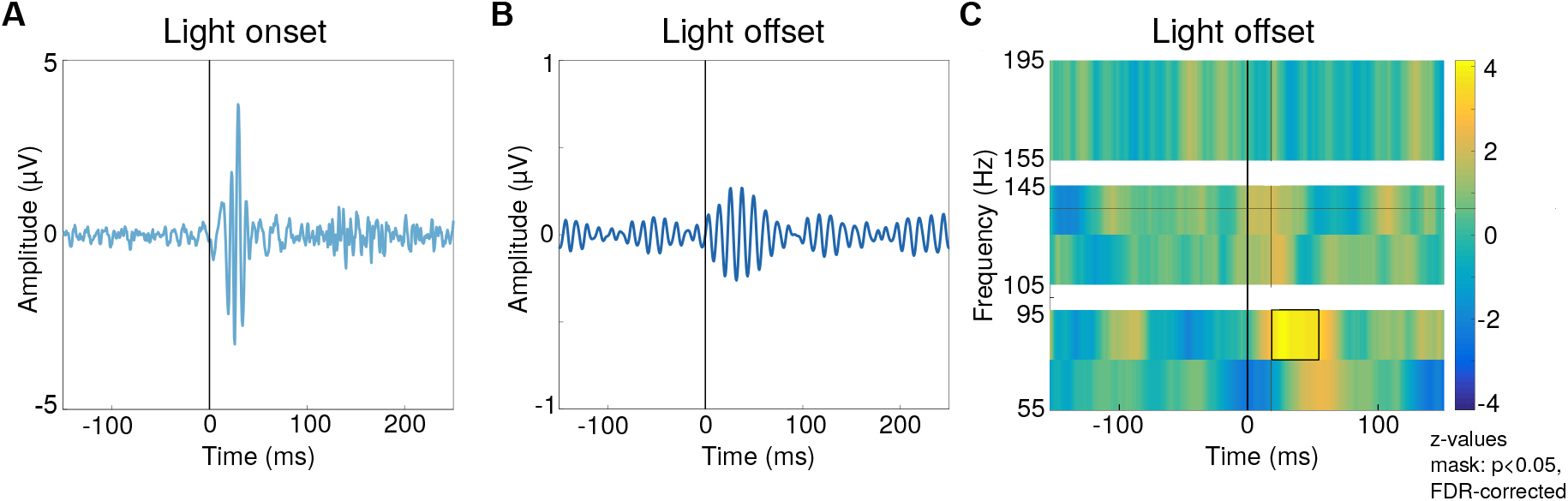
Retinal oscillatory potentials. **A** Oscillatory potential after light onset. The data is highpass filtered at 55 Hz and averaged across subjects. **B** The oscillatory potential after light offset. Data is bandpass filtered from 75 to 95 Hz and averaged across subjects. **C** Intertrial coherence for light offset, tested against baseline across subjects (Wilcoxon rank sum test), black box marks significant area (*p* < 0.05, corrected for multiple comparisons by controlling the false discovery rate [FDR]).

**Figure 3:**
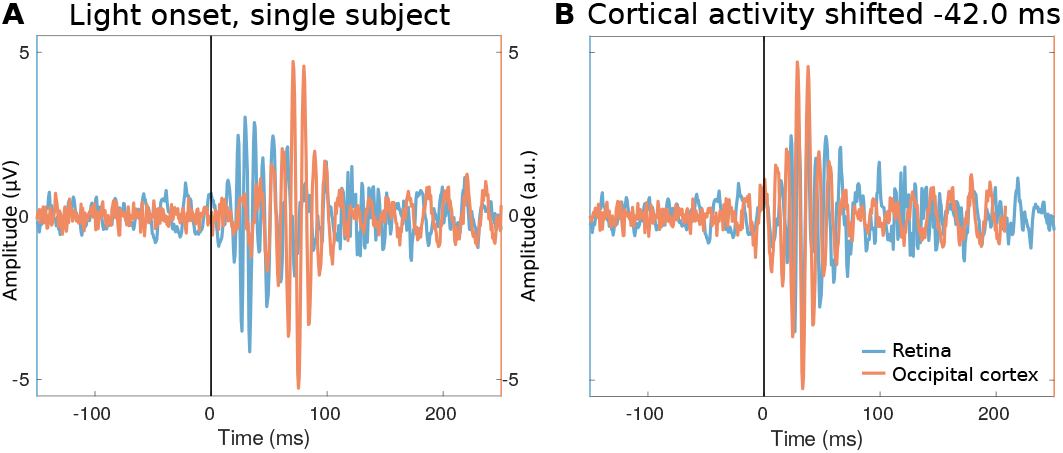
Retinal and cortical high frequency activity in a representative subject. **A** The retinal oscillatory potential (light blue) is plotted together with the cortical high frequency activity of the occipital voxel with the maximal activity. All activity is high-pass filtered at 55 Hz. **B** The same activity as in A, but aligned in time by shifting the occipital activity such that the maxima of retinal and cortical activity line up (resulting in a shift of −42.0 ms for the cortical activity in this subject).

Cortical activity was measured using a 306-channel MEG system (Neuromag TRIUX, Elekta Instruments, Stockholm, Sweden) in a magnetically shielded room. To evaluate high frequency activity in the MEG data, Hilbert amplitudes of five frequency bands (55–75, 75–95, 105–125, 125–145, and 155–195 Hz) were computed in source space, using an linearly constrained minimum variance (LCMV) beamformer and individual head models. In response to light onset, all but the highest frequency band show significant increases in Hilbert amplitude relative to baseline (Wilcoxon rank sum test across subjects). Figure 4A shows this broadband response (the black boxes indicate significant time periods, *p* < 0.001, corrected for multiple comparisons by controlling the false discovery rate [FDR]) and suggests that the activity in higher frequency bands occurs earlier than changes in lower frequencies. This gamma band activity spans occipital regions, including V1 as well as upstream visual regions (Figure 4B, masked for *p* < 0.001, FDR-corrected). In contrast, responses following light offset comprised a narrowband response of 75–95 Hz localized to occipital cortex (Figure 4C and D, masked for *p* < 0.001, FDR-corrected), which is the same frequency band as for the retinal response to light offset.

**Figure 4:**
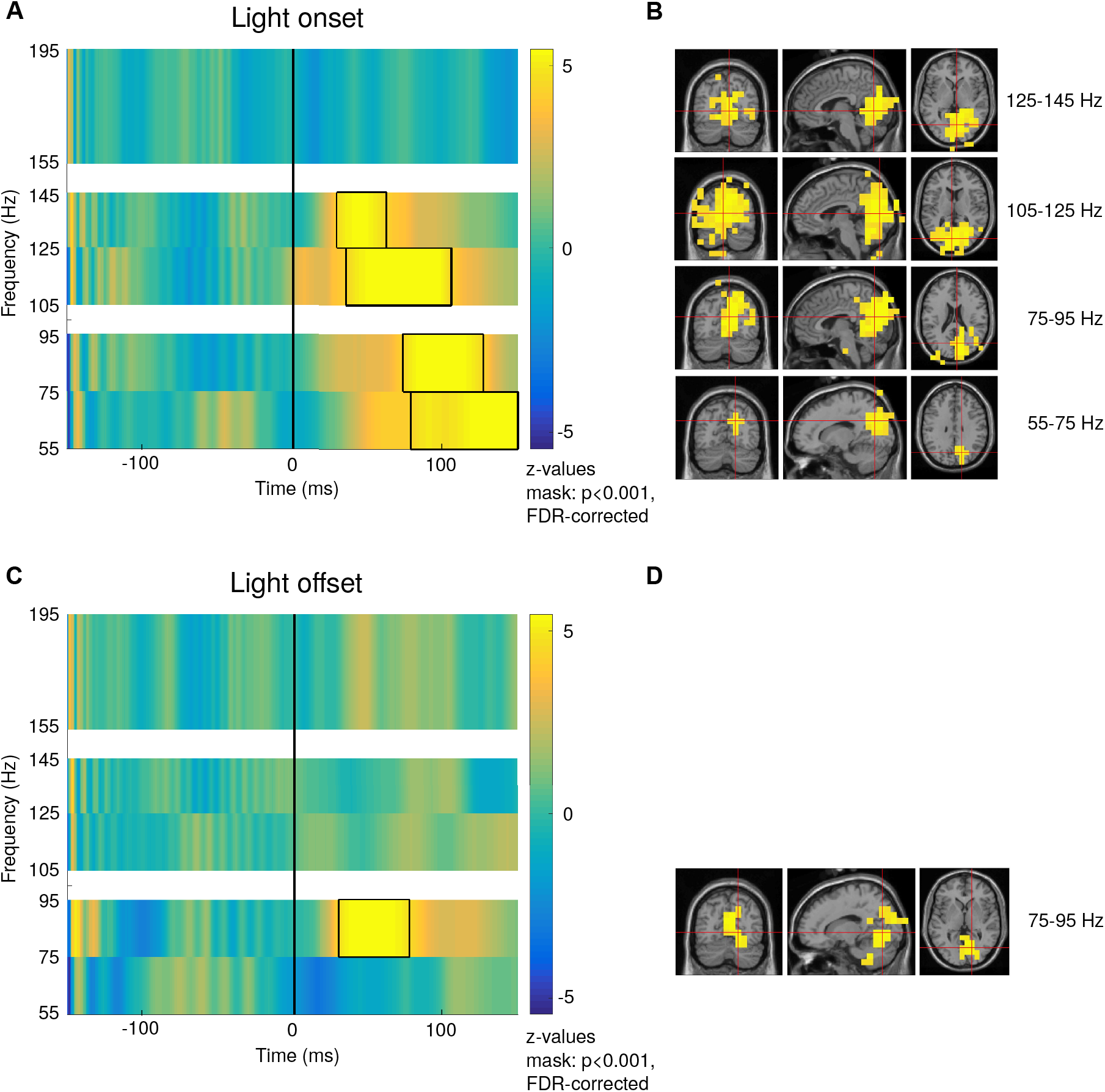
Occipital high frequency activity. Both light onset and light offset evoked cortical high frequency activity. Hilbert amplitudes in different frequency bands are tested against baseline across subjects with the Wilcoxon rank sum test. Depicted for each frequency band is the time courses of the maximum voxel. **A** Occipital high frequency activity following light onset. Black boxes indicate time periods with *p* < 0.001 (FDR-corrected). **B** Voxels with significant activity in the different frequency bands following light onset, activity is masked for p < 0.001. The red cross hairs mark the maximum voxels. **C** High frequency activity following light offset in visual cortex. **D** Localization of flash offset activity, the cross hairs mark the maximum voxel.

It has recently been shown in rats that the flash-induced retinal oscillatory potential is transferred via the optic nerve to the occipital cortex (Todorov et al., 2016). Figure 3 illustrates how similar the retinal and cortical bursts of high frequency activity are for a representative subject. To investigate and compare retinal and cortical oscillatory potentials in this study, we calculated the intertrial coherence (ITC) for those frequency bands that revealed significant power increases (see above). Figure 5A shows the ITC time course in response to light onset for the ERG (depicted in pale blue, median and interquartile range (IQR) across subjects) and for occipital cortex (pale red), where the cortical ITC time course is computed as the across-subjects median of individual occipital maximum ITC. It is evident that the retinal oscillatory potential after light onset is followed by phase-consistent activity in the cortex (individual subject activity is depicted in Supplementary Figure S1). This pattern is most obvious for the higher frequency bands: in the 105–125 Hz band, the cortical ITC peak follows after 51.0 ms (median difference between retinal and cortical peak across subjects, *n* = 9, since the retinal onset ITC peak was not identifiable for one subject). In the 125–145 Hz band, cortical activity follows the retinal ITC peak after 32.0ms (*n* = 9). As illustrated in Figure 5B, light offset led to a comparable pattern: an increase in trial-wise phase consistency in the ERG (depicted in dark blue) is followed by increased ITC in occipital cortex after 21.0ms (*n* = 7), which is shown in dark red (also compare Table 1). An estimation of functional connectivity between the retinal and cortical time-series shows significantly increased Granger causality from retina to cortex compared to the opposite direction (cluster permutation test, *p* < 0.01). This effect is most pronounced for frequencies from 110.0–136.0 Hz following light onset, and for a narrower frequency range (75.3–88.0 Hz) after light offset (Figure 6A). When reversing the time series, the connectivity pattern reverses as well (Figure 6B), as expected for non-spurious results (Haufe et al., 2013; Winkler et al., 2015).

**Figure 5:**
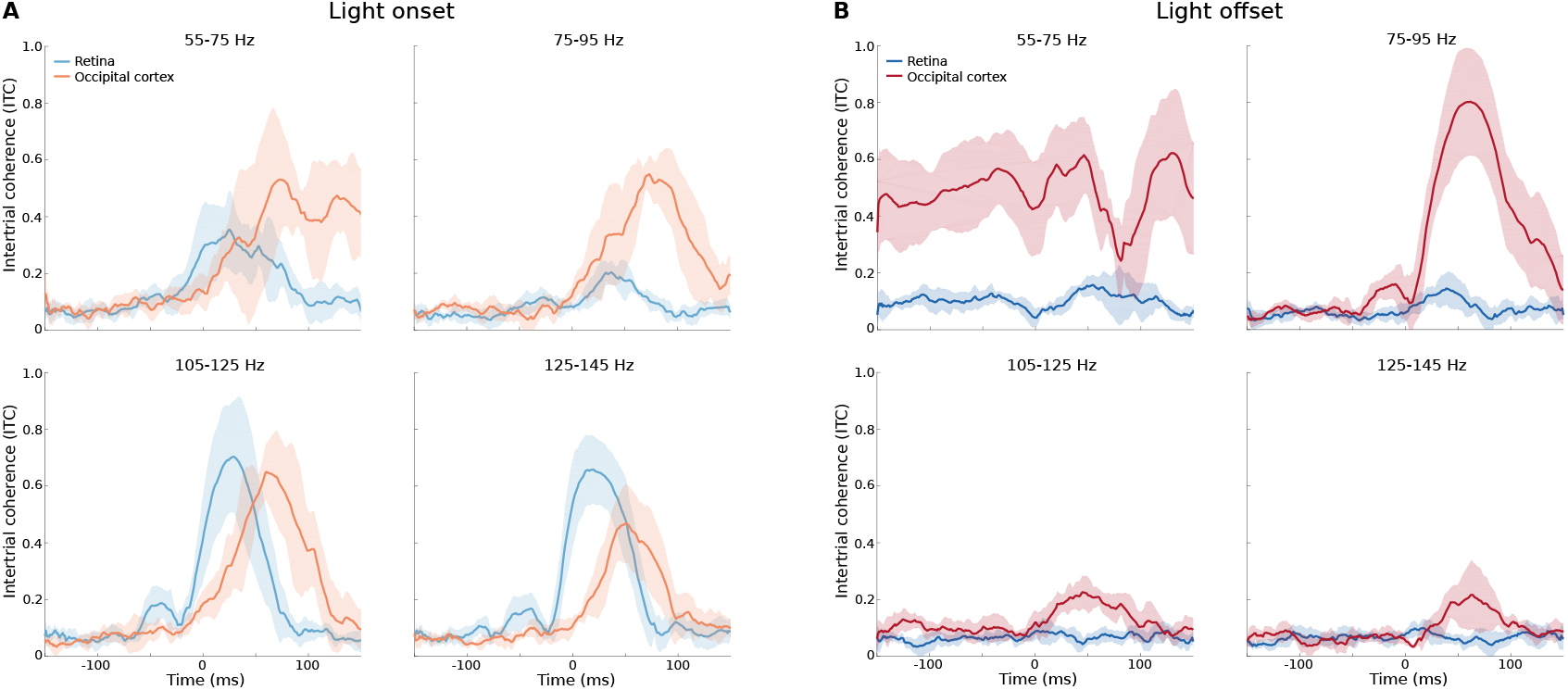
Retinal and cortical intertrial coherence. Shown are median time courses across subjects, shaded areas represent the interquartile range. **A** Time courses of ITC following light onset for the frequency bands with significant activity (cf. Figure 4A). Retinal ITC courses are depicted in pale blue, the cortical ITC in pale red. Cortical ITC time courses represent the median across subjects’ individual maximum ITC activity in occipital cortex. **B** ITC time course in response to light offset for all frequency bands. Note that only the 75-95 Hz frequency band showed significant activity for the offset response (cf. Figure 2C and 4C). The retinal response is shown in dark blue, the median response across individual occipital maximum voxels is shown in dark red.

**Figure 6:**
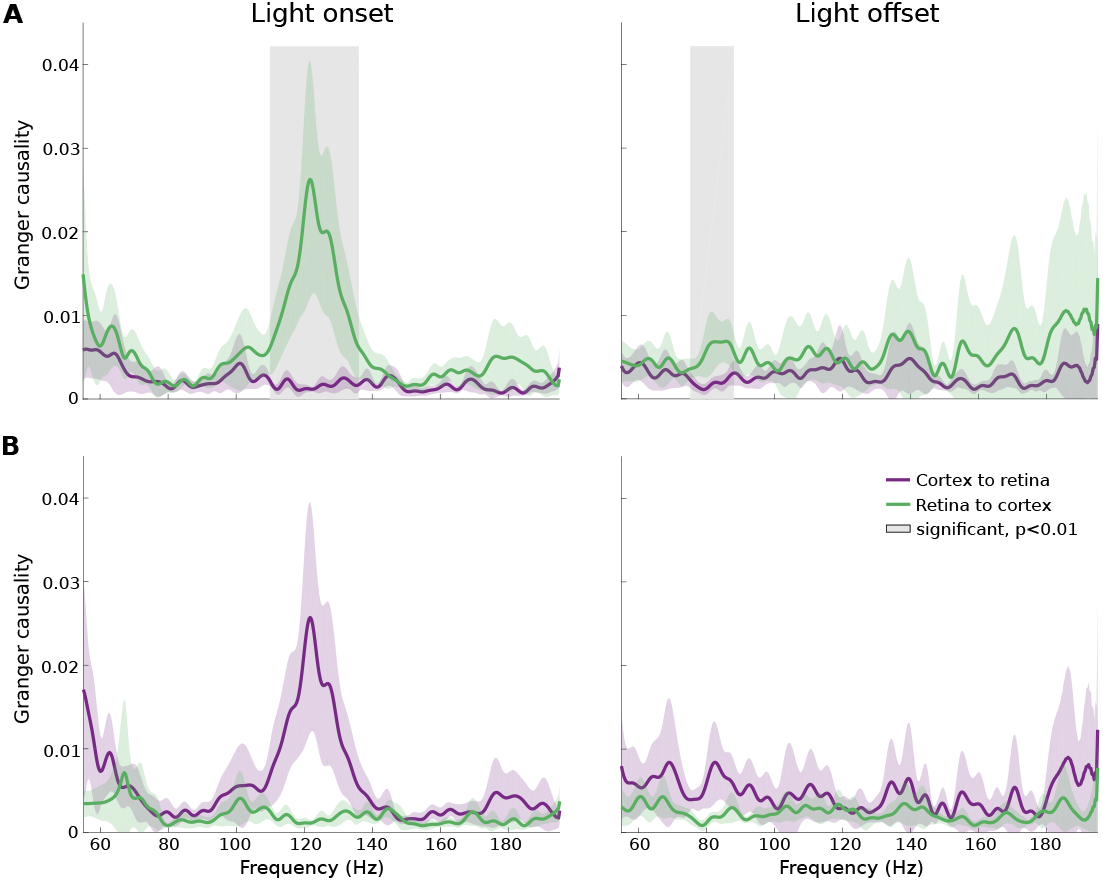
Granger connectivity patterns between retina and cortex. Shown are the Granger causality spectra for frequencies between 55 and 200 Hz. Granger causality estimated for the directionality of “retina to cortex” are depicted in green, while estimates for “cortex to retina” are depicted in purple. Light colored areas represent the standard deviation. **A** Granger spectra for light onset (left) and light offset (right). Grey shaded areas represent contiguous frequencies showing a difference in directionality. **B** Granger spectra for time-reversed data to rule out spurious effects.

**Table 1:** Intertrial coherence peak latencies. Retinal and cortical peak times for the ITC of different frequency bands for both light onset and offset. Shown are median peak times for frequency bands with significant activity (cf. Figures 2 and 4), and the interquartile range (IQR), the interval between the 25^*th*^ and 75^*th*^ percentiles. Specified as well are the number of identifiable peak times per condition (subjects, *n*). The last column shows the results from a Bayesian T-Test of light onset peak latencies of the different frequency bands against the 75-95 Hz light offset peak latency. Boldfaced values indicate a Bayes Factor larger than 3.0, which is considered at least moderate evidence (Jeffreys, 1961; cf. Method details). The last section of the table shows the propagation times (median across subjects, IQR, and number of subjects), defined as the difference between the cortical and retinal ITC peak time, including the results from a Bayesian T-Test examinating the difference between light onset and light offset propagation time.

|  | Light onset |  |  | Light offset |  |  | Bayes factor |
| --- | --- | --- | --- | --- | --- | --- | --- |
|  | median | IQR | *n* | median | IQR | *n* |  |
| Retina |  |  |  |  |  |  |  |
| 55-75 Hz | 31.5 ms | 21.50 | 8 |  |  |  | 0.568 |
| 75-95 Hz | 34.0 ms | 12.00 | 9 | 34.0 ms | 9.75 | 7 | 0.500 |
| 105-125 Hz | 27.0 ms | 3.75 | 9 |  |  |  | **9.931** |
| 125-145 Hz | 27.0 ms | 6.00 | 10 |  |  |  | **3.461** |
| Occipital cortex |  |  |  |  |  |  |  |
| 55-75 Hz | 69.0 ms | 20.50 | 9 |  |  |  | 1.196 |
| 75-95 Hz | 71.5 ms | 22.50 | 10 | 57.0 ms | 18.00 | 10 | **5.911** |
| 105-125 Hz | 77.0 ms | 27.00 | 10 |  |  |  | **4.082** |
| 125-145 Hz | 58.0 ms | 15.50 | 9 |  |  |  | 2.246 |
| Propagation times retina – cortex |  |  |  |  |  |  |  |
| 55-75 Hz | 40.0 ms | 15.00 | 7 |  |  |  | 1.465 |
| 75-95 Hz | 38.0 ms | 32.00 | 9 | 21.0 ms | 17.75 | 7 | 1.698 |
| 105-125 Hz | 51.0 ms | 27.50 | 9 |  |  |  | **4.077** |
| 125-145 Hz | 32.0 ms | 14.25 | 9 |  |  |  | 2.154 |

### High frequency activity: comparing light onset and offset

To assess whether the latencies of retinal and cortical high frequency bursts differ between light onset and offset, ITC peak times were tested across subjects with Bayesian T-Tests. Since light offset was characterized by a narrowband response (75–95Hz), the different onset frequency bands were all tested against this one offset frequency band. The retinal ITC peak latencies (cf. Figure 7) show no support for a difference between light onset and offset for the frequency bands of 55–75 Hz (Bayes Factor: *BF* = 0.591) and 75–95Hz *(BF* = 0.581; cf. Table 1). In the 105–125Hz and 125–145Hz frequency band, the light onset ITC peaks were earlier (both peaking at 27.0 ms on average, *n* = 9 and *n* = 10, respectively) than the 75–95Hz light offset peak (34.0ms, *n* = 7), *BF* = 3.960 and *BF* = 10.62. In the cortex, however, there is support for an earlier offset response peak time at 75–95 Hz (57.0 ms, *n* = 10) than the light onset oscillatory potentials of the 75–95Hz frequency range (71.5ms, *n* = 10), *BF* = 6.451, and the 105–125 Hz frequency range (77.0 ms, *n* = 10), *BF* = 3.397. There is no support for a difference for the other light onset frequency bands of 55–75 Hz (*BF* = 1.556) and 125–145Hz (*BF* = 0.631; cf. Table 1 and Figure 7). Thus, whereas offset high frequency oscillations peak faster than onset responses in the cortex, they seem to peak slower in the retina. This pattern suggests that the narrowband light offset response is transferred faster to cortex than the onset response: the light offset ITC peaks later in the retina than the light onset response for 105–125 Hz, but earlier in the brain. This is supported by the results of testing the propagation time for light onset against that of light offset: in the 105–125 Hz frequency band, there is evidence for a faster transmission of light offset responses (*BF* = 4.077, cf. lower section of Table 1).

**Figure 7:**
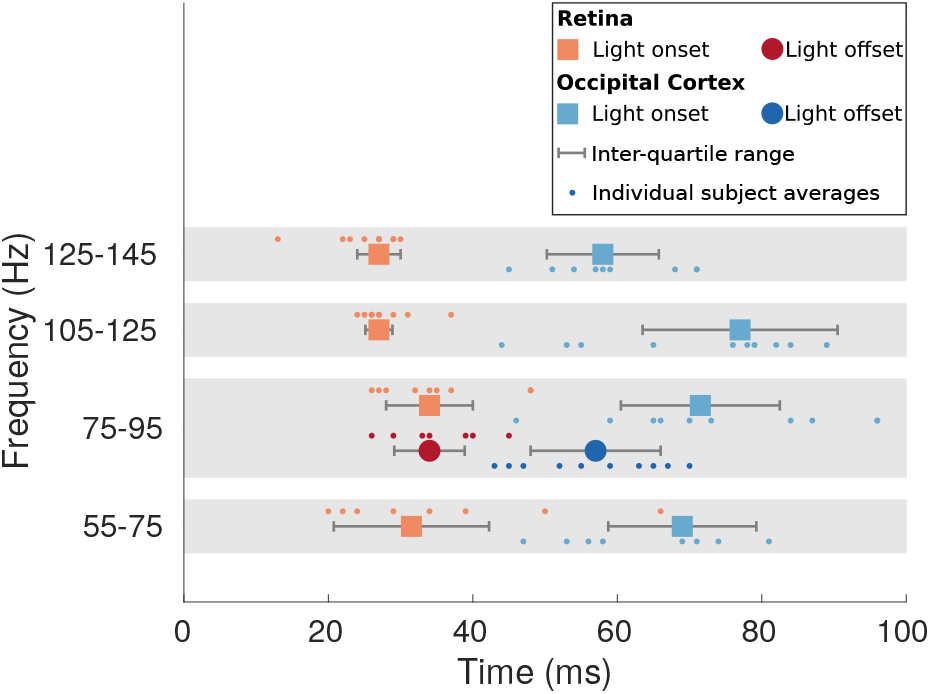
Comparison of intertrial coherence peaks for light onset and offset. The figure shows the median ITC peak latencies across subjects. Red hues represent retinal peak latencies, blue hues refer to cortical responses. The light coloured squares represent the broadband response following light onset (for both retina and cortex), while the dark coloured dots depict the peak latencies of the narrowband light offset response. The errorbars depict the interquartile range (IQR), the interval between the 25^*th*^ and 75^*th*^ percentiles. The smaller dots represent individual subject averages. Note that the 75–95 Hz frequency band is broadened solely for visualization purposes.

## Discussion

We simultaneously recorded retinal and cortical activity to flash onsets and offsets in order to investigate the processing of darks and lights in the human visual system. While numerous studies show advantages for the processing of darks on different levels of the visual system (e.g., Zemon et al., 1988; Yeh et al., 2009; Jin et al., 2008; Jin et al., 2011; Komban et al., 2011; Nichols et al., 2013; Komban et al., 2014), electrophysiological evidence from the human brain is still sparse. We focused on oscillatory activity in the high frequency range, hypothesizing that these high frequency responses would facilitate long-distance coupling between the retina and occipital cortex, as suggested for the visual systems of the cat (Castelo-Branco et al., 1998; Neuenschwander et al., 2002) and rat (Todorov et al., 2016), but also for the human brain (e.g., Fries, 2005; Schoffelen et al., 2005; Fries, 2015).

### Are retinal oscillatory potentials transmitted to cortex?

The present study used retinal and cortical high frequency activity to investigate relative propagation times for darks and lights. This approach was motivated by recent findings that high frequency oscillations enable the functional coupling of retina, thalamus, and visual cortex in mice and rats (Todorov et al., 2016; Saleem et al., 2017; Storchi et al., 2017). Although many studies provide strong evidence that the retinal oscillatory potential propagates through the visual system (Lopez and Sannita, 1997; Castelo-Branco et al., 1998; Sannita et al., 1999; Heinrich and Bach, 2001; Munk and Neuenschwander, 2000; Neuenschwander et al., 2002; Koepsell et al., 2009; Todorov et al., 2016; Saleem et al., 2017; Storchi et al., 2017), there is still some controversy around this topic. Some studies indeed suggest that there is no retinocortical propagation of high frequency activity: Heinrich and Bach (2004) described different peak frequencies in retina and cortex and Molotchnikoff et al. (1975) reported a lack of cortical high frequency activity following flash-stimulation despite a clear retinal response; Doty and Kimura (1963) found a link between retinal and cortical gamma band activity in monkeys but not in cats. Moreover, the nature of a possible propagation is debated as well: Todorov et al. (2016), for example, reported high coherence between the retina, the optic chiasm, and visual cortex in rats; however, they note differing waveform shapes in these three stages of the visual system and therefore argue against a merely passive spread of the oscillatory potential.

Our data show evoked oscillatory activity following light onset and offset in both retina and cortex. This activation comprises similar frequency bands in the retina and in visual cortex: the light onset response is broadband in the retina (55–195 Hz) as well as in cortex (55–145 Hz; whereupon the lack of significant activity in the 155–195 Hz frequency band is presumably due to the low signal-to-noise ratio of such high frequencies in MEG data). Equivalently, the offset response is restricted to the same frequency band (75–97 Hz) in both retina and visual cortex. Furthermore, an analysis of functional connectivity revealed significantly increased Granger causality from retina to cortex in both conditions. Crucially, the frequencies with the most pronounced effects closely map the frequency bands found in the other analyses. Following light onset, there is an increase in Granger causality in a frequency band from 110–136 Hz, coinciding with the two frequency bands that showed the highest ITC values. In the offset response, Granger causality was enhanced in a lower, narrower frequency band (75.3–88Hz), coinciding with the ITC increase in the 75–97Hz frequency band.

The Granger causality results as well as the ITC activity patterns are consistent with the propagation of the retinal oscillatory potential to visual cortex. Furthermore, the difference in propagation time between light onset and offset responses revealed through the ITC analysis suggests involvement from the thalamus, as discussed in more detail below, and thus indicates that the propagation of the oscillatory potential to the visual cortex is not a mere passive spread, corroborating Todorov et al. (2016). Such long-distance gamma coupling has been previously shown in other modalities, e.g., between the spinal cord and motor cortex (Schoffelen et al., 2005), and has been hypothesized to facilitate communication (Fries, 2005; Fries, 2015). In that theoretical model, *communication through coherence* (CTC), the rhythmic synchronization of postsynaptic neuron groups through interneuron networks leads to a rhythmic modulation of input gain for these postsynaptic groups (Fries, 2015). Consequently, only coherent neuronal groups communicate effectively. While the CTC model could explain our data, it interestingly does not conform with the results by Todorov et al. (2016). They recorded the oscillatory potential also from the optic chiasm, which they point out does not possess nerve cells that could generate such a rhythmic synchronization. Todorov et al. (2016) hypothesize instead, that the oscillatory potential might spread along the axon membrane of the optic nerve as a synchronous local field oscillation.

Remarkably, our data also shows the existence of retinal oscillatory activity following light offset. Such activity has before been reported in cats, where a 65–100 Hz high frequency oscillation was recorded after light offset (Kozak, 1971), but not yet in humans. We call this high frequency activity the *dark retinal oscillatory potential (DROP)*.

### Are darks processed faster than lights?

Behavioral studies suggest that darks are processed faster than lights: reactions to dark objects and light decrements are faster and more accurate (Blackwell, 1946; Chubb and Nam, 2000; Buchner and Baumgartner, 2007). At the cortical, thalamic, and retinal levels, evidence for faster processing of darks is mixed (e.g., Lankheet et al., 1998; Chichilnisky and Kalmar, 2002; Gollisch and Meister, 2008; Jin et al., 2008; Yeh et al., 2009), although numerous studies report greater neural resources for the processing of darks (e.g., Balasubramanian and Sterling, 2009). Functional evidence from the human visual system, however, is sparse.

To resolve temporal differences in processing and propagation in the visual system, we compared retinal and cortical ITC peak times for five different high frequency bands ranging from 55–195 Hz.

In the retina, light onset peaks *earlier* than light offset in two frequency bands: 105–125 Hz and 125–145 Hz. For the lower frequency bands, there is no support for a difference concerning light onset and offset peak times. In visual cortex, however, the narrowband 75–95 Hz light offset response clearly peaks faster than the activity in the two predominant frequency bands for light onset, 75–95 Hz and 105–125 Hz.

Taken together, the peak latencies of oscillatory activity in our experiment thus suggest the processing of darks on the cortical level is faster, but slower on the retinal level. Due to the fact that the ERG represents the summed activity of different cell types and due to the controversy regarding the exact origin of the retinal oscillatory potential (Doty and Kimura, 1963; Perlman, 2001; Kenyon et al., 2003; Frishman, 2013), it is difficult to speculate about the precise underlying retinal mechanisms of this finding. However, given that the retinal b-wave may peak *after* the initial responses of visual cortex (Clark et al., 1994; Shigihara et al., 2016), the oscillatory potential may better reflect the timing of the retina’s output stages. On the cortical level, these findings corroborate the hypothesis of faster processing of darks than lights (Komban et al., 2014).

Emerging from the retinal and cortical peak latencies of oscillatory activity, this suggests a shorter time lag between retinal and cortical processing for darks (21.0 ms) than lights (32.0 to 51.0 ms, which replicates the previously reported lag for flash onsets of 48ms [Heinrich and Bach, 2001]). For the 105–125Hz frequency range, statistical testing yields evidence for a difference in propagation times. Based on the fact that the retinal high frequency activity emerges from late stages of processing (Doty and Kimura, 1963; Perlman, 2001; Kenyon et al., 2003; Frishman, 2013), whereas the cortical high frequency response is the first activity arising in visual cortex after visual stimulation, this time lag can be used as a proxy for propagation time in the visual system. Our findings then can be interpreted as showing a faster retinocortical propagation time following light offset compared to light onset, thus pointing beyond a merely cortical effect. This could be explained by faster and more efficient processing of darks by the thalamus, which would be supported by the finding of faster processing for light decrements in the LGN of cats (Jin et al., 2011). Correspondingly, Xing et al. (2010) described a temporal advantage for darks in the thalamic input layer of V1 in macaque monkeys, although they report no time differences in upstream visual areas. While the findings of Jin et al. (2011) are specific to the thalamic pathways of cats, functional magnetic resonance imaging (fMRI) as well as modelling work point to the complexity of the LGN in humans, exhibiting temporal modulations that can originate from LGN properties or cortical feedback loops (for review, see Ghodrati et al., 2017). The hypothesis that thalamus is responsible for the propagation differences we report here could be tested in a follow-up study, exploiting methods that can capture thalamic activity. An alternative explanation for the diverging transmission times could be functional asymmetries extending beyond the retina through to cortex. From an information processing perspective, the lesser informational complexity for darks compared to lights could be another key aspect regarding the propagation time differences. Light contains more complex visual information than darkness, and light onset could evoke more complex visual scene processing, e.g., stereo vision, potentially prolonging thalamic processing.

### Slow retinal potentials

We additionally compared the retinal evoked potentials following light onset and offset. Studies on the generators of these potentials suggest that the d-wave evoked by light offset is analogous to the b-wave evoked by light onset, since both are presumably driven by bipolar cells (Sieving et al., 1994; Perlman, 2001; Frishman, 2013; Vukmanic et al., 2014). In our data, the light offset d-wave peaks faster than the light onset b-wave. However, the fact that the b-wave peaks as late as 79.9 ms (which is even later than the ITC peaks of high frequency activity in cortex [58.0 to 77.0ms]) raises the question of whether this is a valid comparison. Anchoring the d-wave and the light onset a-wave to the respective oscillatory potentials yields further indications for this: the d-wave latency is around 10 ms earlier than the latency of the *dark retinal oscillatory potential*. The same is true for the light onset activity when comparing the a-wave latency to the peak times of the different frequency bands, whereas the b-wave peaks over 45 ms after this high frequency activity. This relation suggests that the d-wave and b-wave might not reflect comparable responses of OFF and ON bipolar cells after all. However, our data suggest that the d-wave may actually be an analogue to the a-wave, as there is no support for a difference between their peak latencies, which would contradict the view that they are generated by different retinal layers (Sieving et al., 1994; Perlman, 2001; Frishman, 2013).

### The role of narrowband and broadband gamma responses in the visual system

As reported above, light onset evoked a broadband high frequency response in the retina and visual cortex, whereas light offset was followed by a narrowband response in the 75–95 Hz range. This dissociation of different bandwidths contributes to the debate about the functional relevance of narrowband and broadband high frequency activity. Narrowband oscillatory activity in the visual system is well known to arise in response to stationary or moving grating stimuli (e.g., Adjamian et al., 2004; Hoogenboom et al., 2006; Muthukumaraswamy et al., 2010) and has also been described with focused attention (Vidal et al., 2006). However, there is a debate about the origin as well as functional implication of such narrowband responses, for example, about the question whether they have general relevance to visual stimuli (Brunet et al., 2014) or whether they are specific to artificial stimuli such as gratings (Hermes et al., 2014; Hermes et al., 2015). In our study, the narrowband gamma response was elicited specifically by light offset. This finding suggests that gratings might be triggering similar responses of the visual pathway as light offset.

A recent paper showed narrowband gamma oscillations in the visual system of mice: Saleem et al. (2017) report that visual broadband and narrowband activity is not correlated and demonstrate that the narrowband gamma response is inherited from thalamus. They propose a model with two different channels for information transfer: the narrowband gamma mediating thalamocortical communication and the broadband gamma effectuating corticocortical interactions. In the present study, however, we show that both narrowband and broadband gamma are indeed transmitted from retina to cortex. We contend that they comprise different levels of informational complexity, as discussed above.

## Conclusions

In summary, our results strengthen findings of faster processing for darks in visual cortex and thereby deliver a possible explanation for any behavioral advantages of darks over lights. On the retinal level, we did not find faster processing of light decrements. Instead, light increments seem to be processed faster than light decrements. This study furthermore shows that both retinal high frequency activity following light onset as well as the *dark retinal oscillatory potential* following light offset get transmitted to visual cortex. Based on this finding, our data can also be interpreted to show faster propagation times for darks, possibly due to faster thalamic processing. Moreover, we show that high frequency activity in response to light onset comprises a broad range of frequencies, whereas the response to light offset evokes a narrowband oscillation in the range of 75–95 Hz in both retina and cortex. This could reflect the higher informational value in lights compared to darks, which could also account for faster thalamic processing.

## Acknowledgements

We thank Christopher Bailey, Kousik Sarathy Sridharan, and Andreas Højlund for their assistance in data collection and Tzvetan Popov and Ursula Lommen for their help with a pilot recording. We thank Mads Jensen for his advice on statistical analyses. Further, we thank Juan Vidal for valuable discussions about this study. This work was supported by the Zukunftskolleg of the University of Konstanz, ERA-Net NEURON via the Bundesministerium für Bildung und Forschung (BMBF grant 01EW1307), and the European Research Council (Starting Grant 640448).

## Materials and methods

### Participant details

10 healthy participants (four female, average age 34.1 years; *s.d*. = 6.31) took part in the study. 6 participants were contact lens wearers, since experience showed that they usually tolerate the ERG electrode very well. Contact lens wearers did not wear their lenses during the experiment. All participants provided written informed consent and the study was approved by the Videnskabsetiske Komitéer for Region Midtjylland, Komite II (Ethical Committees of Central Denmark Region, Committee II) and carried out in accordance with the Declaration of Helsinki.

## Method details

### Stimuli and experimental setup

The experimental stimuli were full field light flashes which were presented using the Presentation software (Neurobehavioral Systems, Inc., Berkeley, CA). The white flashes had a duration of 480ms and were followed by a black screen which was shown for a random time interval between 2000 and 2500 ms. A total of 250 flashes was shown and the experiment lasted approximately 12min. The flashes were projected onto a screen inside the MEG chamber using a ProPixx projector (VPixx Technologies Inc., Saint-Bruno, Canada) with a 120Hz refresh rate and symmetric rise and fall times < 1 ms (cf. Supplementary Figure S2). Participants were seated in an upright position, the projection screen was at 70 cm distance from the subjects. The flashes were as full-field as possible subtending the central 28.01° (vertical extent) and 47.77° (horizontal extent) of the visual field and had a brightness of 280 cd/m^2^.

### Data acquisition

MEG data was recorded using a 306-channel MEG system (102 magnetometers and 204 gradiometers, Neuromag TRIUX, Elekta Instruments, Stockholm, Sweden) in a magnetically shielded room. Data was sampled at 5 kHz with a recording bandwidth of 0.1–1650 Hz. Bilateral ERG was recorded using disposable Dawson-Trick-Litzkow (DTL) fiber electrodes. Additionally, horizontal and vertical electrooculogram (HEOG and VEOG) were recorded using a bipolar montage. The ERG electrodes were referenced to the ipsilateral HEOG. Prior to data acquisition, the head position indicator (HPI) coils and three fiducial points (left and right periauricular points and nasion) were digitized using a Polhemus Fastrak 3D scanner (Polhemus, Colchester, VT, USA) for later coregistration with the structural magnetic resonance image (MRI) of the subjects. The on- and offsets of the flashes were recorded with a photodiode during the whole experiment.

### Statistical analysis

Analysis of MEG and ERG data was conducted using the open-source toolboxes Field-Trip (Oostenveld et al., 2010) and NUTMEG (Dalal et al., 2004; Dalal et al., 2011) for MATLAB, Bayesian analyses were fitted in JASP (JASP Team, 2018). Epochs of light onsets and offsets were identified using the photodiode traces. Trials with eye-movements were rejected based on the HEOG and VEOG activity. Subsequently, trials including muscle artifacts or MEG channel jumps were excluded as well, leaving on average 183.4 trials (*std* = 24.06) per subject and condition. The data was downsampled to 1000 Hz.

### ERG data

For ERG data analysis, only data from the left ERG was used. Data was baseline corrected and detrended and the epochs were then averaged with respect to light onset and offset. The peak latencies for the retinal potentials (a-, b- and d-wave) were identified on the averaged time series for every subject. A paired samples Wilcoxon signed rank test was conducted on the b-wave and d-wave measurements, as well as on the a-wave and d-wave peaks. To obtain the oscillatory potentials after light onset, ERG data was highpass-filtered at 55 Hz (Hanning windowed FIR filter, onepass-zerophase, 6Hz transition width). For light offset, data was highpass-filtered at 75 Hz and lowpass-filtered at 95 Hz using the same filter definitions.

### MEG data

For MEG data analysis, only the 102 magnetometers were used. Boundary element head models with three layers (brain, skull, scalp) were constructed for every subject based on the individual structural MRI using OpenMEEG (Gramfort et al., 2010; Gramfort et al., 2011). The source grid spanning the whole brain had a resolution of 10 mm. Sources were reconstructed using the linearly constrained minimum variance (LCMV) beamformer (Van Veen et al., 1997) with normalized weights (Van Veen et al., 1997; Sekihara and Nagarajan, 2008). The covariance matrices passed to the beam-former were computed based on the Minimum Covariance Determinant estimator, providing a robust covariance matrix estimate. The beamforming approach was combined with the Hilbert transform to acquire source space Hilbert amplitude and phase for five frequency bands: 55–75, 75–95, 105–125, 125–145, and 155–195 Hz. To generate these frequency bands, separate high- and lowpass filters were adopted (Hanning windowed FIR filter, onepass-zerophase, 6Hz transition width). For every frequency band and condition, a spatial filter was constructed as described above, and the single trials were projected through the filter to yield virtual electrodes at every grid point. Subsequently, the time courses of the virtual electrodes were Hilbert transformed, providing amplitude estimates for every frequency band. Intertrial coherence (ITC) for timepoint t and frequency bands f was computed based on the Hilbert phase estimates as a measure of phase synchrony across trials, using the following formula: 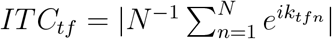, with *N* being the total number of trials and *e^ik^* representing the complex polar representation of the phase angle *k* on the *n^th^* trial (cf. Cohen, 2014). To allow for comparison of retinal and cortical high frequency activity, the ERG data was filtered and Hilbert transformed in the same manner. To compute Granger causality, the epoched MEG data was source reconstructed using an LCMV beamformer. Spectral content was then computed for both the ERG and beamformed MEG data using the fast Fourier transform (FFT). Frequency smoothing of 5 Hz was applied using a discrete prolate Slepian sequence (DPSS) multitaper approach. Spectrally resolved Granger analysis following Bastos and Schoffelen (2016) was computed between the occipital peak voxel activity and the retinal time series for each subject. Significant differences in Granger causality for direction were determined using cluster permutation tests (Maris and Oostenveld, 2007). 10000 permutations were computed to establish statistical significance while controlling for mutliple comparisons, using a non-parametric approach to determine the clusters. To control for spurious effects in connectivity, Granger causality was also computed and reported for the time-reversed data (Haufe et al., 2013; Winkler et al., 2015).

### Statistical testing

For both MEG and ERG data, the Hilbert amplitude time courses were normalized against the distribution of baseline time points using the Wilcoxon rank sum test for every frequency band and subject. The z-values obtained from this step were tested against the baseline distribution across subjects with the Wilcoxon rank sum test. The derived statistics were corrected for multiple comparisons by controlling the false discovery rate (FDR). ITC peaks were identified for every subject; in source space, the maximal ITC was searched among all occipital voxels based on the Automated Anatomical Labeling (AAL) atlas (Tzourio-Mazoyer et al., 2002) for every subject. Due to a polyphasic response, the retinal light offset ITC for one participant was smoothed by lowpass filtering the ITC time series (cutoff: 40 Hz) to identify a clear peak latency. Light onset and offset ITC peak times were then tested using a Bayesian Paired Samples T-Test for both retinal and cortical activity. Due to missing values, propagation times were tested using a Bayesian Independent Samples T-Test. All Bayesian tests used a normally distributed prior, centered around 0, with r = 0.707. Results are reported as Bayes factors (BF), which express to which extent the data supports a model under the alternative hypothesis over a model under the null hypothesis (Lee and Wagenmaker, 2014). Bayes factors are interpretable as odds ratios, however, they are frequently reported according to an evidence categorization by Jeffreys (1961), which regards BF > 3 as moderated evidence and BF > 10 as strong evidence for the model that represents the alternative hypothesis (Lee and Wagenmaker, 2014).

## Data availability

The data that support the findings of this study are available from the corresponding author upon reasonable request and the completion of a data processing agree ment(https://medarbejdere.au.dk/en/informationsecurity/data-protection/general-information/data-processing-agreements/) in accordance with Aarhus University policies.

## Code availability

The code used to analyze the data and generate the figures of this study is available under https://github.com/britta-wstnr/retina_cortex_darks_lights.

