## Supplementary information for "Faster than the brain’s speed of light: Retinocortical interactions differ in high frequency activity when processing darks and lights"

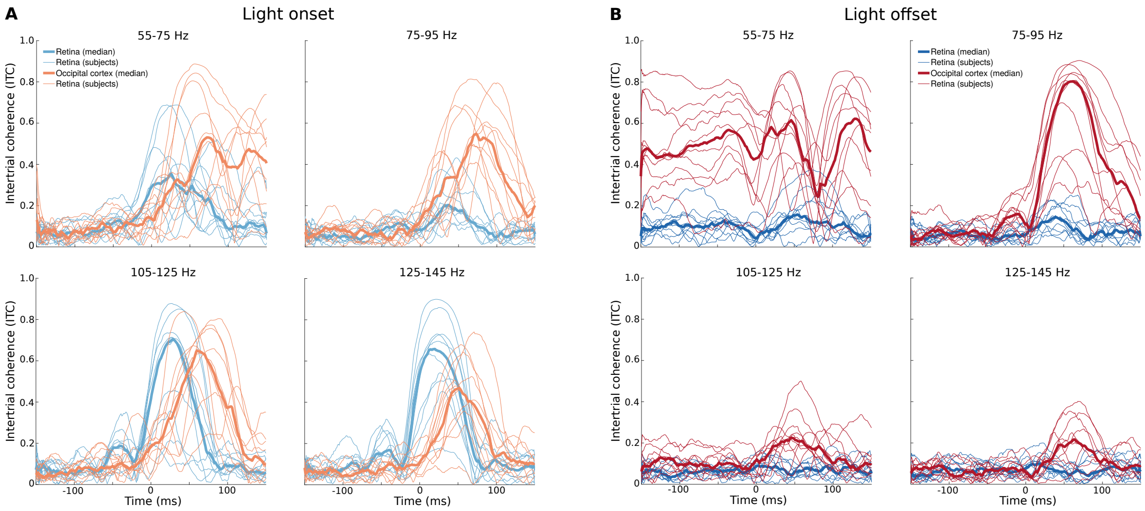


**Figure S1. Retinal and cortical intertrial coherence: Individual subjects.** Shown are median time courses across subjects (thick line) and individual subject averages across trials (thin lines). **A** Time courses of intertrial coherence (ITC) following light onset for the frequency bands with significant activity. Retinal ITC courses are depicted in pale blue, the cortical ITC in pale red. Cortical ITC time courses represent the subjects' individual maximum ITC activity in occipital cortex. **B** ITC time course in response to light offset for all frequency bands. Note that only the 75 Hz to 95 Hz frequency band showed significant activity for the offset response. The retinal response is shown in dark blue, the response in the individual occipital maximum voxels is shown in dark red.

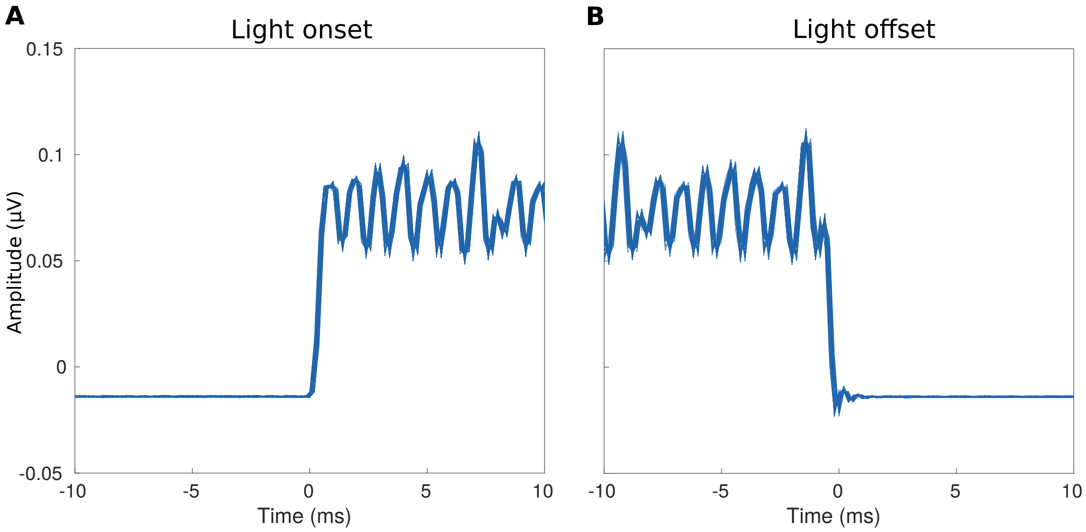


**Figure S2. Rise and fall times of the visual stimulus.** Depicted are photodiode measurements of the projection screen at light onset and light offest. 100 trials are shown overlaid (not averaged), demonstrating their extremely high consistency. Rise and fall times are <1 ms. **A** Stimulus onset (light onset) photodiode traces, 100 trials. **B** Stimulus offset (light offset) photodiode traces, 100 trials.
